# Cross-talk between individual phenol soluble modulins in *S. aureus* biofilm formation

**DOI:** 10.1101/2020.04.01.020610

**Authors:** Masihuz Zaman, Maria Andreasen

## Abstract

The infective ability of the opportunistic pathogen *Staphylococcus aureus* is associated with biofilm mediated resistance to host immune response and even disinfectants and indeed *S. aureus* is recognized as the most frequent cause of biofilm associated infections. Phenol-soluble modulin (PSM) peptides serve various roles in pathogenicity while also comprising the structural scaffold of *S. aureus* biofilms through self-assembly into functional amyloids, but the role of the individual PSMs during biofilm formation remains poorly understood and the molecular pathways of PSM self-assembly have proved challenging to identify. Here, we show a high degree of cooperation between individual PSMs during the formation of functional amyloids in biofilm formation. The fast aggregating PSMα3 initiates the aggregation by forming unstable aggregates capable of seeding the formation of aggregates by other PSM peptides into the formation of stable amyloid structures. Using chemical kinetics along with spectroscopic techniques we dissect the molecular mechanism of aggregation of the individual peptides to show that PSMα1, PSMα3 and PSMβ1 display secondary nucleation whereas βPSM2 aggregates through primary nucleation and elongation. Our findings suggest that the various PSMs have evolved to ensure fast and efficient biofilm formation through cooperation between individual peptides.

## INTRODUCTION

Aggregated proteins in the form of functional amyloids are widespread in Nature [1]. In humans, functional amyloids assist in immunity, reproduction and hormone secretion [2]. However, in various bacterial strains they provide structural stability as the major protein component of the self-produced polymeric matrix in biofilms [3-6]. Functional amyloids increase the bacteria’s ability towards a variety of environmental insults, increasing their persistence in the host as well as promoting resistance to antimicrobial drugs and the immune system [7-9]. The well-studied curli machinery in *Escherichia coli* [3], Fap system in *Pseudomonas fluorescens* [4], TasA system in *Bacillus subtilis* [5], along with phenol-soluble modulins (PSMs) in *Staphylococcus aureus* [6] are some of the major bacterial functional amyloid systems that have been reported so far.

For *S. aureus* biofilm formation PSMs have been recognized as a crucial factor. In their soluble monomeric form they hinder host immune response by recruiting, activating and lysing human neutrophils while also promoting biofilm dissociation [6]. However, self-assembly of PSMs into amyloid fibrils fortify the biofilm matrix to resist disassembly by mechanical stress and matrix degrading enzymes [10]. The genes encoding the core family of PSMs peptides are highly conserved and located in *psm*α operon (PSMα1-PSMα4), *psm*β operon (PSMβ1 and PSMβ2) and the δ-toxin is encoded within the coding sequence of RNAIII [11]. High expression of PSMαs, ∼20 residues in length, increases virulence potential of methicillin-resistant *S. aureus* [12]. Moreover, PSMα3 the most cytotoxic and lytic PSM, enhances its toxicity to human cells upon fibrillation [13]. Despite lower concentrations, the larger PSMβs, ∼44 residues in length, seem to have the most pronounced impact on biofilm structuring [14]. Despite the formation of functional amyloids in *S. aureus* by PSMs, many questions remain about the intrinsic molecular mechanism by which they self-assemble and what molecular events triggers the formation of fibrillar structure from their monomeric precursor peptide. Here, we apply a combination of chemical kinetic studies along with biophysical techniques to explore the relative importance of different microscopic steps involved in the mechanism of fibril formation of PSMs peptides.

## RESULTS

### Chemical kinetics reveal different aggregation mechanisms for different PSMs

To investigate the dominating mechanism of aggregation for the individual PSMs we used chemical kinetics to analyze the aggregation of all the seven individual PSM peptides under quiescent conditions. Recently, kinetic models of protein aggregation [15, 16] have been effectively applied to numerous model systems in biomolecular self-assembly [17-19]. Through these models, the aggregation kinetics ascertain the rates and reaction orders of the underlying molecular events, allowing for the determination of the dominating molecular mechanism of formation of new aggregates. Aggregation kinetics of all seven PSMs peptides (PSMα1-4, PSMβ1-2 and δ-toxin) was monitored using Thioflavin T (ThT) fluorescence intensity [20]. For PSMα1, PSMα3, PSMβ1 and PSMβ2 reproducible aggregation curves were observed, Fig 1a-c and 1g, while for the rest of the PSM peptides (PSMα2, PSMα4 and δ-toxin) no reproducible aggregation was observed, Fig. S1a-c. An increase in ThT fluorescence is observed for PSMα4 although this was not sigmoidal in shape and also not reproducible, Fig. S1b. The timescale for the completion of aggregation differs significantly between the PSM ranging from ∼1 h for PSMα3 and up to ∼70 h for PSMα1. Furthermore, the aggregation of PSMβ1 was carried out at concentrations of µg/mL compared to concentrations at mg/mL for the other PSM peptides since at higher concentrations of PSMβ1 the lag-time during aggregation becomes monomer independent suggesting a saturation effect, Fig S1d.

**Fig. 1:**
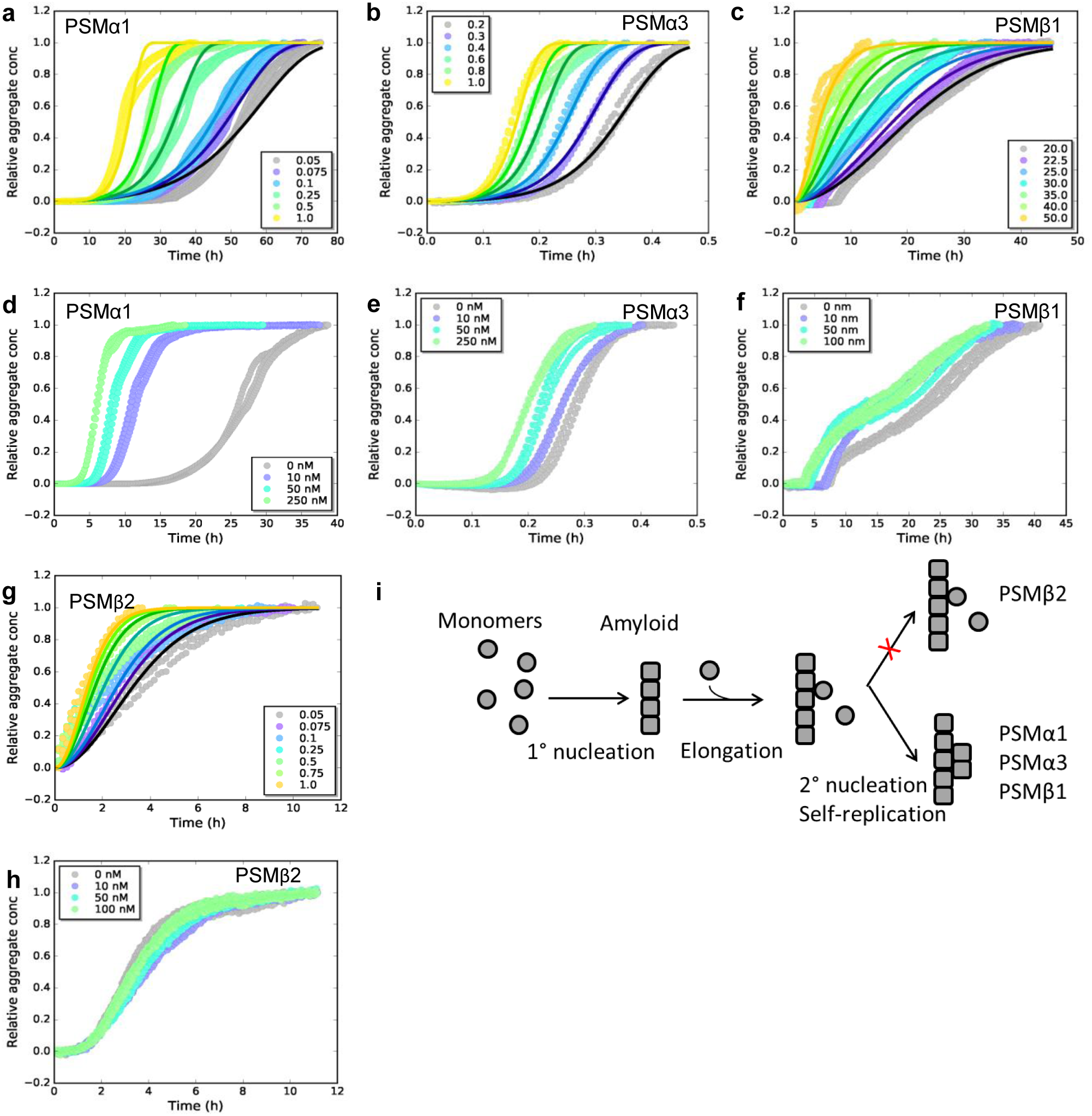
Experimental ThT kinetic data for the aggregation of PSMs peptides from monomeric samples. (a) Aggregation of PSMα1 (0.05 mg/mL to 1.0 mg/mL). The data are well fitted by a secondary nucleation model. (b) Aggregation of PSMα3 (0.2 mg/mL to 1.0 mg/mL). The data are well fitted by a secondary nucleation model. (c) Aggregation of PSMβ1 (20 µg/mL to 50 µg/mL). The data are well fitted by a secondary nucleation model. (d) Aggregation of PSMα1 in the presence and absence of low concentrations of preformed seeds, at a monomer concentration of 0.5 mg/mL and seed concentration of 0 to 250 nM monomer equivalents. Significant effects of added seeds on the rate of aggregation were observed. (e) Aggregation of PSMα3 in the presence and absence of low concentrations of preformed seeds, at a monomer concentration of 0.4 mg/mL and seed concentration of 0 to 250 nM monomer equivalents. Significant effects of added seeds on the rate of aggregation were observed but lower than αPSM1. (f) Aggregation of PSMβ1 in the presence and absence of low concentration of preformed seeds, at a monomer concentration of 0.025 mg/mL and seed concentration of 0 to 100 nM monomer equivalents. Significant effects of added seeds on the rate of aggregation were observed but lower than αPSM1. (g) Aggregation of PSMβ2 (0.05 mg/mL to 1.0 mg/mL). The data are well fitted by a nucleation-elongation model. (h) Aggregation of PSMβ2 in the presence and absence of low concentration of preformed seeds, at a monomer concentration of 0.5 mg/mL and seed concentration of 0 to 100 nM monomer equivalents. No significant effects of added seeds on the rate of aggregation are evident. (i) A schematic illustration of the microscopic steps involved in PSM aggregation. Monomers of PSMβ2 nucleate through primary nucleation with rate constant k_n_ and the aggregates grow by addition of monomers to the fibril end with rate constant k_+_. In addition to primary nucleation and elongation monomers of PSMα1, PSMα3 and PSMβ1 nucleate through secondary nucleation on the surface of an already existing aggregate with secondary rate constant k_2._ All kinetic experiments were carried out in triplicates. Parameters from the data fitting are summarized in Table 1.

To elucidate the dominating aggregation mechanism of PSMα1, PSMα3, PSMβ1 and PSMβ2, kinetic data were globally fitted at all monomeric concentrations concurrently by kinetic equations using the Amylofit interface (http://www.amylofit.ch.cam.ac.uk/fit) [21]. High quality global fits were achieved for all four peptides assuming a secondary nucleation mechanism for PSMα1, PSMα3 and PSMβ1, and a primary nucleation and elongation mechanism for PSMβ2. The presence of a single dominating aggregation mechanism for all four peptides is seen in the linear correlation between the half-time and the initial monomer concentration [21], Fig. S2a-d. In this simple nucleation-elongation (or linear self-assembly) model, the protein monomers form an initial nucleus with rate constant (*k*_*n*_) and reaction order (*n*_*c*_) which grow by elongation through the addition of monomers to the fibrils ends with rate constant (*k*_*+*_). The secondary nucleation model additionally involves nucleus formation catalyzed by existing aggregates. In this model system, *k* act as a combined parameter that controls the proliferation through secondary pathways with secondary process rate constant (*k*_*2*_) and secondary pathway reaction order (*n*_*2*_) with respect to monomer [21]. Secondary nucleation dominated aggregation mechanisms have previously been reported for disease related amyloid fibrils, for example Aβ peptides [17, 18], insulin [22], α-synuclein [23, 24] and islet amyloid poly peptide [25] whereas the nucleation-elongation model has previously been linked to functional amyloids from *E. coli* and *Pseudomonas* [26].

**Table 1:** Kinetic parameters obtained from fitting of data in Figure 1 using the web server AmyloFit. n_c_ and n_2_ are the reaction order of the primary and secondary nucleation process respectively, k_n_ and k_2_ are rate constants for the primary and secondary nucleation process and k_+_ is the rate constant for the elongation of existing fibrils.

| Parameters | PSMα1 | PSMα3 | PSMβ1 | PSMβ2 |
| --- | --- | --- | --- | --- |
| Dominating mechanism | Secondary nucleation | Secondary nucleation | Secondary nucleation | Nucleation-elongation |
| MRE | 0.00311 | 0.00138 | 0.00333 | 0.00475 |
| $k_+k_n$ ($M^{-n_c}h^{-2}$) | 0.0000698 | 275 | $1.86 \times 10^{18}$ | 48.8 |
| $n_c$ (-) | 0.00000784 | 0.6 | 3.92 | 0.572 |
| $k_+k_2$ ($M^{-n_c}h^{-2}$) | 129 | $5.17 \times 10^6$ | $4.23 \times 10^3$ | - |
| $n_2$ (-) | 0.00166 | 0.123 | 0.2 | - |

In order to confirm the dominating mechanism of aggregation of the four peptides based on chemical kinetics, seeded aggregation in a regime of very low seed concentrations to monomeric concentration was conducted. This type of experiments delivers a direct means of probing the ability of fibrils to self-replicate [24, 27]. The presence of preformed fibril seeds can accelerate the aggregation process by two different mechanisms, namely elongation and surface catalyzed secondary nucleation. The presence of low amounts of seeds eliminates the rate limiting step of primary nucleation when secondary nucleation is present but no changes in the kinetics will be observed when only primary nucleation and elongation is present as the low amounts of seed do not eliminate the need for more nuclei to be formed before the elongation process dominates. Indeed a decrease in the lag phase was observed with increasing seed concentration (up to 0.1% of the total protein mass present at the start of reaction) for PSMα1, PSMα3 and PSMβ1, Fig 1d-f and 1h supporting the observation that secondary nucleation is the dominating molecular mechanism for the formation of new aggregates of PSMα1, PSMα3 and PSMβ1 peptide [28]. Despite the very fast kinetics of PSMα3, the reduction in lag phase was significant and can be clearly observed, Fig. 1e. The aggregation kinetics of PSMβ2 was not affected by the presence of low amounts of seeds confirming the lack of self-replication processes in the form of surface catalyzed secondary nucleation, Fig. 1h. The general mechanism underlying formation of new aggregates from monomers of the PSM peptides from both primary and secondary pathways is shown in Fig. 1i.

### Elongation rates differ by a factor of 1000 between fastest and slowest PSM

The relative contributions of elongation rate constant (*k*_*+*_) were investigated in presence of high concentration of preformed fibril seeds. The global fitting of the kinetic data yields a product of the elongation rate constant and the primary nucleation rate constant (*k*_*n*_*k*_*+*_), however, in the presence of high amounts of preformed seeds, the intrinsic nucleation process becomes negligible, and hence the aggregation under this type of experiments is only dependent on elongation of the aggregates [28]. The initial increase in aggregate mass was measured through linear fits to the early points of the aggregation process, Fig. S3a, S3c, S3e and S3g [29, 30]. The estimated elongation rate constant for PSMα1 and PSMβ2 were found to be 0.2 mM/h^2^ and 0.5 mM/h^2^ respectively, differing by a factor of ∼2, which is insignificant. Contrary to PSMα1 and PSMβ2 the estimated elongation rate constant for PSMβ1, which aggregates at very low monomeric concentrations, was found to be 0.2 µM/h^2^ and hence a factor 1000 smaller than for PSMα1 and PSMβ2. In contrast, the elongation rate constant of PSMα3 was found to be 16.6 mM/h^2^ exceeding the values of PSMα1 and PSMβ2 ∼80 and ∼35 fold, respectively. The stronger effect of elongation of PSMα3 in comparison with other three peptides suggests that interactions with the fibrils increases the importance of PSMα3 fibrils in the assembly reactions compared with the assembly of free monomers of other peptides in the solution.

### Secondary structure analysis confirm α-helical structure of PSMα3 and β-sheet structure for other PSMs

The changes in secondary structure of the peptides following aggregation was monitored using synchrotron radiation circular dichroism (SRCD) spectroscopy and Fourier transform infrared (FTIR) spectroscopy. The CD spectra of monomers of all PSMs peptides prior to aggregation all show double minima at 208 and 222 nm indicative of α-helical structure, Fig S4a. Upon aggregation the CD spectrum of the peptides changes displaying a single minimum at approximately 218 nm for PSMα1 and PSMα4, and at 220 nm for PSMβ1 and PSMβ2 indicative of β-sheet rich structure, Fig 2a. Despite the lack of sigmoidal aggregation curves for PSMα4 changes in the CD spectrum upon incubation was observed. This indicates a transition from α-helical structure to a structure with increased β-sheet content upon aggregation and is consistent with data previously published [4, 5]. The CD spectrum of aggregated PSMα3 is still displaying a double minimum with minima shifted to 208 nm and 228 nm indicative of α-helical structure being present in the aggregates although this helical structure is different form that observed in the monomeric peptide. This observation is consistent with the reported cross-α-helical structure of PSMα3 aggregates [13]. To further probe the contribution of the individual structural components to the CD spectra each spectrum was deconvoluted using the analysis programs Selecon3, Contin and CDSSTR in the DichroWeb server [31, 32], Fig. 2b and Table S1. Indeed the major structural contribution to the SRCD spectrum for PSMα3 aggregates is α-helical (∼70%). For PSMα1 and PSMα4 the major structural components are β-sheet (33% and 35% respectively) and unordered structure (39 % and 34 % respectively). Despite the single minima indicative of predominantly β-sheet structure observed for PSMβ1 and PSMβ2 the major structural components are α-helix (35 % and 40 % respectively) and unordered structure (33% and 35 % respectively) with less contribution from β-sheet structure (24 % and16 % respectively). No structural changes were observed for PSMα2 and δ-toxin which upon incubation still displayed spectra characteristic of α-helix, Fig. S4b. This is consistent with the lack of aggregation seen for these peptides by ThT fluorescence.

**Figure 2:**
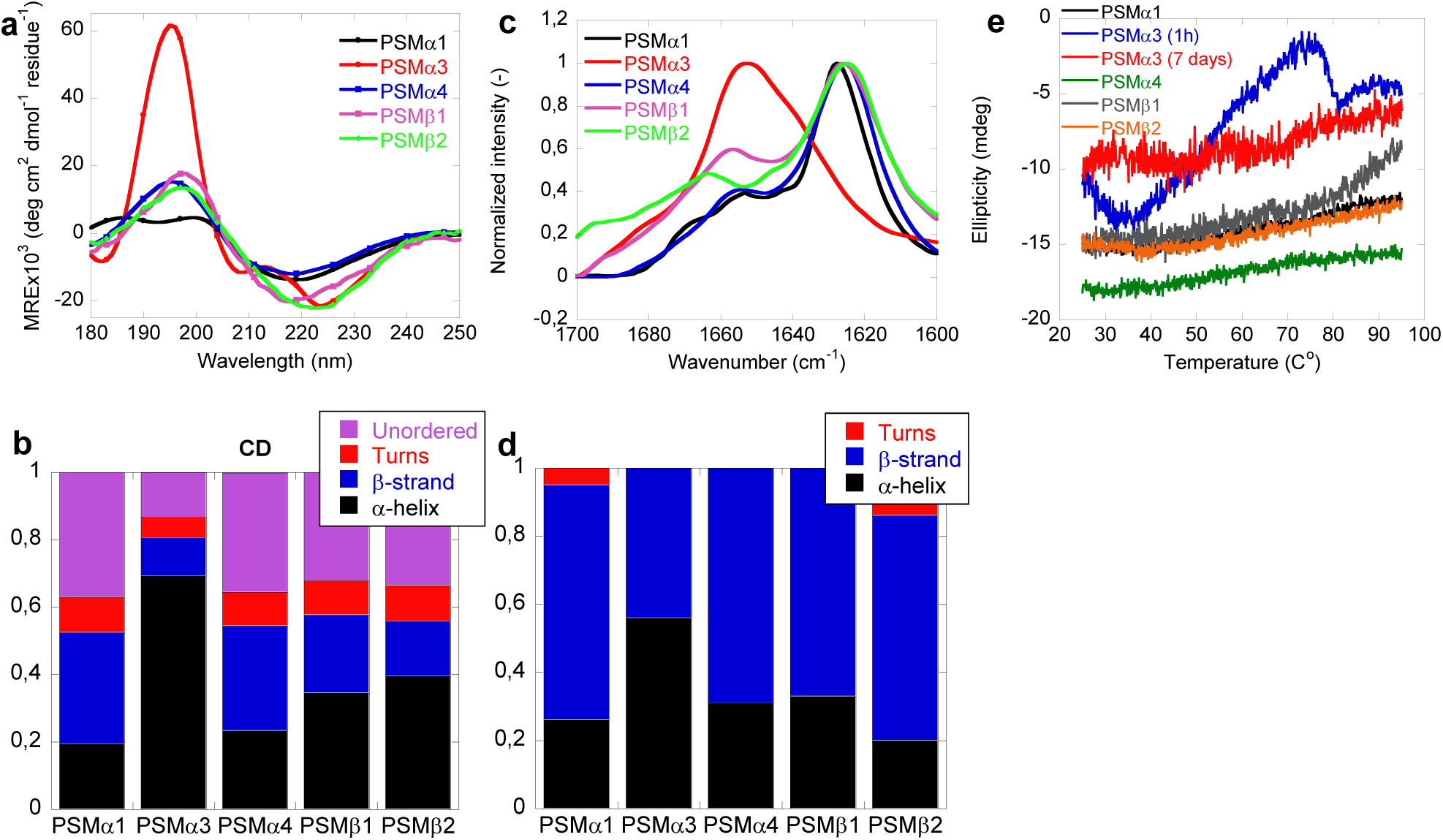
Structural comparison of fibrils formed by different PSMs variants. (a) Synchrotron radiation (SR) Far UV-CD spectra of PSMα1, PSMα3, PSMα4, PSMβ1 and PSMβ2 fibrils recorded after seven days of incubated samples except for PSMα3 which is recorded after one hour of incubated samples. (b) Deconvolution of the SR-CD spectra of fibrils of PSM variants into the individual structural components. (c) Attenuated total internal reflection Fourier transform infrared (ATR-FTIR) spectroscopy of the amide I’ region (1600-1700 cm^-1^) of fibrils of PSMs variants. PSMα1, PSMα4, PSMβ1 and PSMβ2 show a peak at 1625 cm^-1^ corresponding to rigid amyloid fibrils. In contrast, PSMα3 shows main peaks at and 1654 cm^-1^, with the latter indicating on more disordered fibrils. (d) Deconvolution of the FTIR spectra of fibrils of the PSM variants into the individual structural components. (e) CD thermal scans from 20 to 95°C of PSMα1, PSMα3 (1h), PSMα4, PSMβ1 and PSMβ2 fibrils.

Consistent with the CD data, the FTIR spectra of PSMα1, PSMα4, PSMβ1 and PSMβ2 were found to be very similar with a well-defined intense peak at ∼1625 cm^-1^ indicative of amyloid β-sheet and a minor shoulder at ∼1667 cm^-1^ indicative of β-turns [33-35], Fig. 2c. The secondary structure composition of the fibrils was estimated using deconvolution of the spectra followed by conventional fitting program and summarized in Table S2, Fig 3b and FigS5a-e. In good agreement with the CD data and previous reports the FTIR spectra of PSMα3 aggregates shows significantly higher content of α-helical structure relative to other PSMs peptide fibrils as shown by a more intense band in the spectrum of at 1655 cm^-1^, indicative of α-helical structure [36].

**Figure 3:**
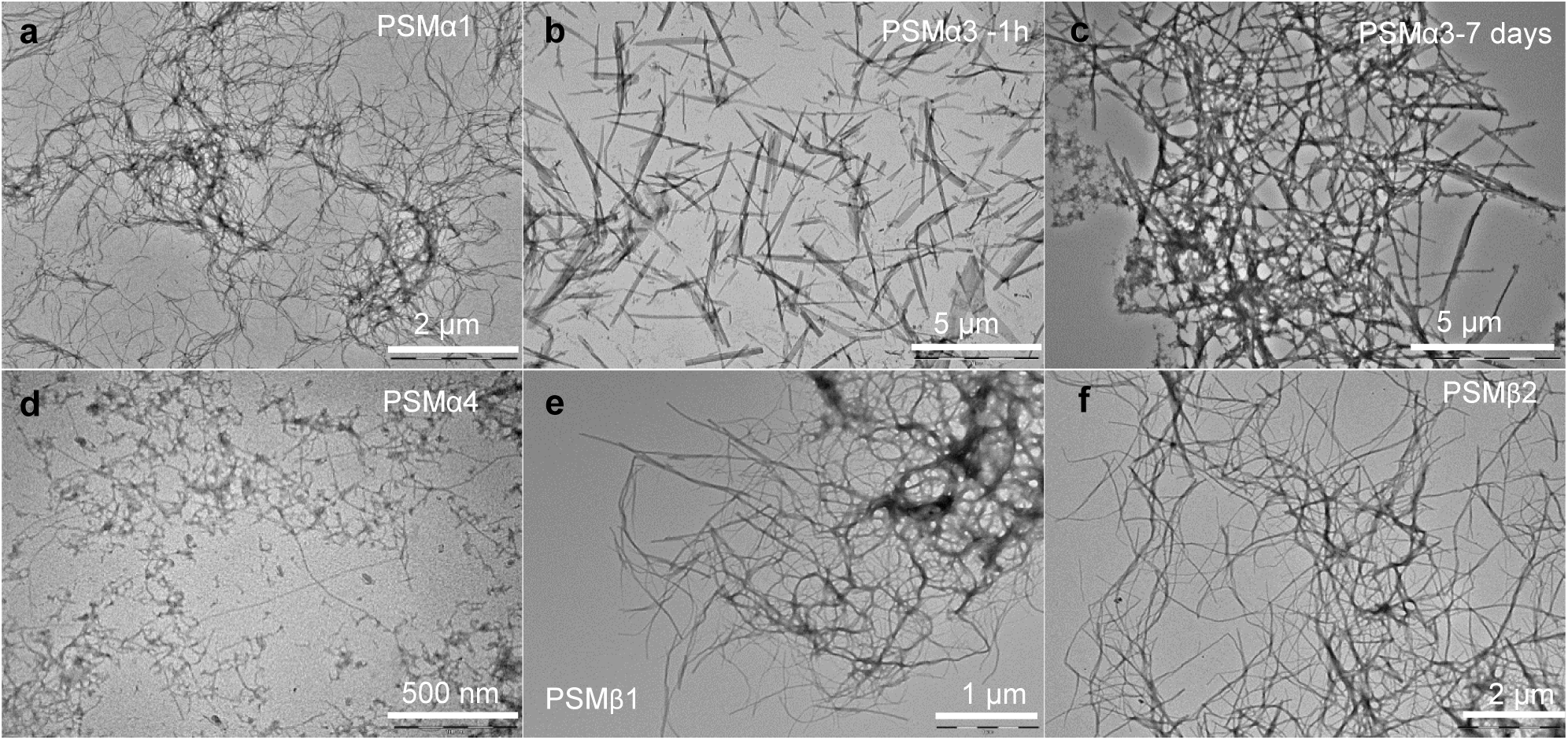
Morphology of aggregates of PSMs peptides. Transmission electron microscopic image of the end state of reaction for samples initially composed of (a) PSMα1 fibrils, (b) PSMα3 fibrils after 1h of incubation, (c) PSMα3 fibrils after 7 days of incubation, (d) PSMα4 fibrils, (e) PSMβ1 fibrils and (f) PSMβ2 fibrils. Please note that scale bar changes.

### Freshly formed PSMα3 aggregates are unstable but becomes stable through lateral association during maturation

The stability of the PSM aggregates was tested using CD spectroscopy. Fig 2e shows the CD signal at 220 nm for fibrils (PSMα1, PSMα3, PSMα4 and PSMβ2) from 25°C to 95°C. All peptide fibrils spectra except PSMα3 indicate thermally stable β-sheet structure. Even at 95°C, there is no indication of the loss of β-sheet structure of PSMα1, PSMα4 and PSMβ2, as judged by the stable negative peak at 220 nm. However, freshly formed (1 h old) αPSM3 fibrils are thermally unstable as a loss of structure is seen above 50°C which is consistent with previous studies of fragments of PSMα3 [37]. Interestingly the lateral association of aggregates of PSMα3 upon further incubation (7 days, see Fig. 3) renders the fibrils thermally stable and no changes in structure is seen upon heating to 95 °C indicating that the lateral association of the aggregates stabilizes the structure. The stability of the aggregates towards chemical denaturants was tested using urea, Fig. S4c-d. Again aggregates of PSMα3 are the only ones susceptible towards disassembly (5M-8M urea) while no apparent effect is observed for PSMα1, PSMα4, PSMβ1 and PSMβ2 fibrils.

The morphological features of aggregates were examined using transmission electron microscopy (TEM). After seven days of incubation, PSMα1 formed stretches of entangled fibrils, Fig. 3a. In addition, bulky dense aggregate surrounded by a network of fibrils were observed. In contrast, PSMα3 incubated for 1h generated short and unbranched fibrils, Fig.3b, which, upon two-day incubation, associates laterally to form stacks, Fig. S6c, and further associate to form entangles networks of fibrils after seven days of incubation, Fig. 3c. Aggregates of PSMβ2 showed entangled fibrils marginally thicker and more dispersed than PSMα1 and PSMα3, Fig. 3f. Aggregates of PSMβ1 also formed entangled networks of fibrils, Fig. 3e. No aggregated species could be observed for PSMα2 and δ-toxin, Fig. S6a-b, consistent with the lack of aggregation as seen by the lack of increase in ThT fluorescence upon incubation. Interestingly, PSMα4 which lacked reproducible ThT kinetics but displayed β-sheet structure using CD and FTIR spectroscopy display very thin fibrils visible at higher magnification with some distribution of spherical aggregates, Fig. 3d. Overall, these data are in good agreement with the recorded kinetics and structural data.

### PSMα1 display promiscuous cross-seeding while other PSMs display selective cross-seeding abilities

The interplay between individual PSM peptides during formation of functional amyloids was investigated using cross-seeding experiments where the ability of aggregates of one PSM peptide to seed the aggregation of the other PSM peptides was tested. Cross-seeding experiments using 20% preformed fibril seeds of PSMα1 and monomers of the rest of the PSM peptides were performed. It can be seen that the aggregation of all the other PSM peptides (PSMα2-4, PSMβ1-2, and δ-toxin) is indeed accelerated by the presence of preformed fibrils seeds of PSMα1, Fig.4a, indicating that PSMα1 seeds not only effectively promoted self-aggregation through both elongation and secondary nucleation but also promoted cross-seeding between all PSMs peptides. Unlike PSMα1 the fast aggregating PSMα3 seeds are only capable of promoting self-aggregation and accelerating PSMα1 aggregation, Fig.4b and Fig. S7. The lag phase of PSMα2, PSMα4, PSMβ1 & δ-toxin did not change upon addition of preformed PSMα3 seeds indicating that PSMα1 is more promiscuous in interaction with the other PSM peptides during aggregation. Interestingly the lag phase of PSMβ2 enhanced dramatically in presence of PSMα3 seeds, Fig.4b.

Like PSMα3 PSMβ1 is only capable of accelerating the aggregation of PSMα1, Fig 4c. Further, the effect of cross seeding other PSM peptides with PSMβ2 seeds was investigated. Pre-formed seeds of PSMβ2 were able to accelerate the aggregation of PSMα1 and PSMβ1 while also inducing aggregation of PSMα2 and δ-toxin, Fig. 4d. None of the other PSM peptides were able to induce aggregation of these two PSM peptides which also do not aggregate on their own under conditions tested here. Furthermore, seeds of PSMβ2 caused insignificant changes in the fibrillations kinetics of PSMα3 and PSMα4.

**Figure 4:**
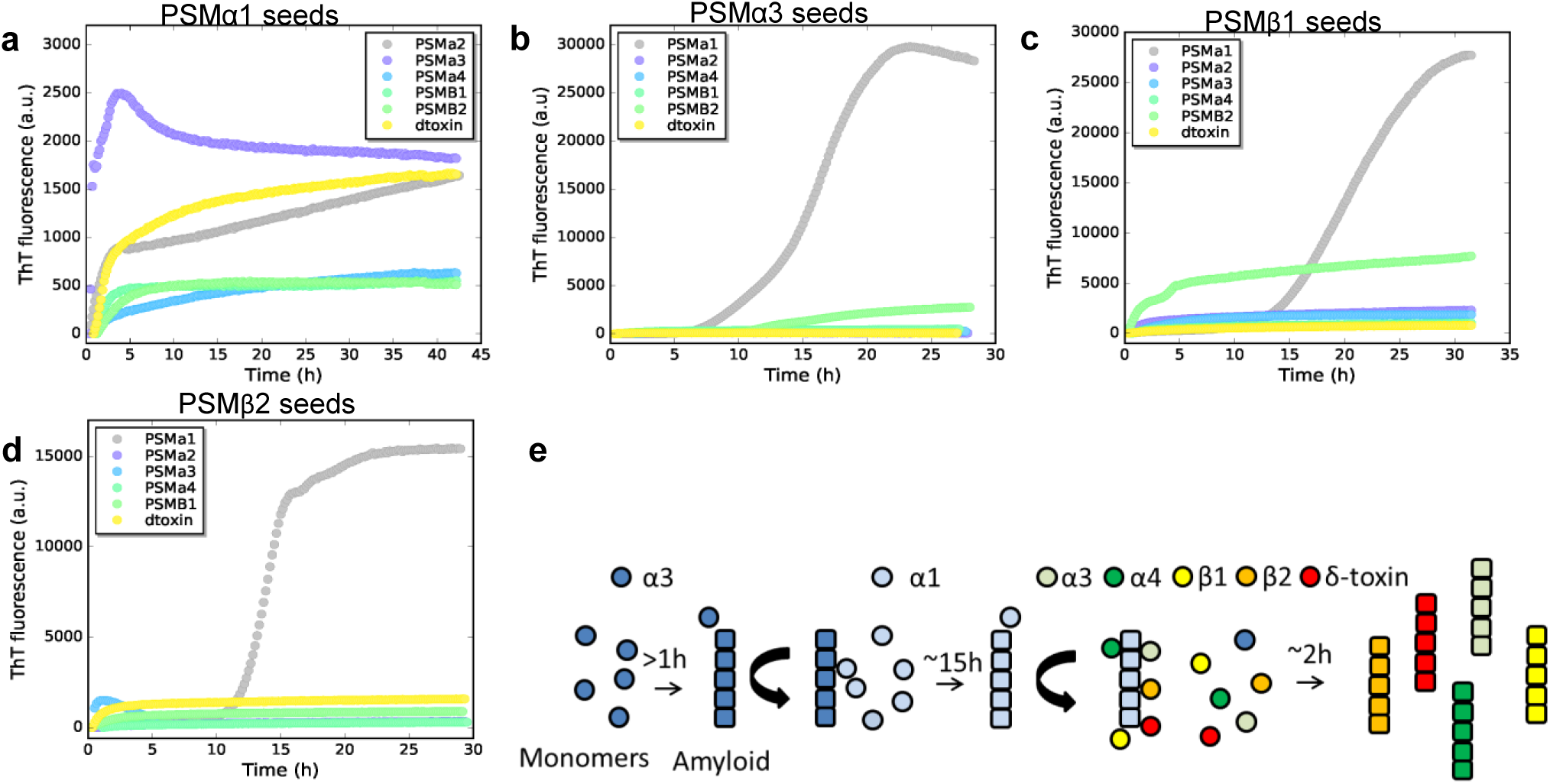
Cross seeding PSMs variant. (a) Aggregation of PSMα2, PSMα3, PSMα4, PSMβ1, PSMβ2 and δ-toxin (0.25 mg/mL) in the presence of 20 % preformed PSMα1 seeds. (b) Aggregation of PSMα1, PSMα2, PSMα4, PSMβ1, PSMβ2 and δ-toxin (0.25 mg/mL) in the presence of 20 % preformed PSMα3 seeds. (c) Aggregation of PSMα1, PSMα2, PSMα3, PSMα4, PSMβ2 and δ-toxin (0.25 mg/mL) in the presence of 20% preformed PSMβ1 seeds. (d) Aggregation of PSMα1, PSMα2, PSMα3, PSMα4, PSMβ1 and δ-toxin (0.25 mg/mL) in the presence of 20 % preformed PSMβ2 seeds. (e) Schematic representation of the cross seeding interactions between the PSM variants during biofilm formation.

Based on the cross-seeding analysis it is clear that the PSM peptides display selectivity in the interaction with preformed aggregates which cannot be explained simply by the sequence similarities with in the PSM α- or β-group indicating an intricate interplay between the various PSM peptides during biofilm formation. It is also clear that PSMα1 is the most promiscuous of the PSM peptides as aggregation of PSMα1 is accelerated by all the other PSM peptides that aggregate while also being able to accelerate the kinetics of aggregation of all the other PSM peptide even the ones that do not aggregate on their own. We therefor suggest a model describing the delicate interplay between individual PSM peptides in the formation of biofilm where the fast aggregating PSMα3 initiates the aggregation by forming unstable fibrils. These fibrils can then accelerate the aggregation of PSMα1 which forms stable fibrils capable of accelerating the aggregation of the majority of the remaining PSM peptides (PSMα2, PSMα3, PSMβ1, PSMβ2 and δ-toxin), Fig. 4e. In this way the fast kinetics of PSMα3 may act as a catalyst for the whole system of aggregation of PSMs peptides during biofilm formation.

## Discussion

PSMs peptides are major determinants and play an important and diverse role in the biofilm matrix in *S. aureus* [11]. Previous studies have shown that PSMs from *S. aureus* form functional amyloids that contribute to biofilm integrity and provide resistance to disruption, which is critical to the virulence of medical device-associated infections [6, 38]. However, there have been no efforts to date to establish a general picture for the self-assembly of PSMs peptides that brings together all the species in the aggregation cascade. In the present study, we have conducted a combination of detailed kinetic analysis with structural and morphological analysis to gain insight into the molecular and mechanistic steps, to determine how functional amyloid of PSMs, the biofilm determinant of *Staphylococcus aureus*, form and grow. This study involves studying separately the different process involved in the aggregation reaction i.e., initial nucleation steps, growth of fibrils and their amplification.

Earlier computational analysis of PSMs sequences (smallest staphylococcus toxins) already suggested that the peptides of individual families might display differential self-assembly properties [38]. We first determined the rates of the various microscopic steps (concentration dependent) associated with the aggregation of PSMs peptides. Our results have shown that under quiescent conditions, a dominant contribution to the formation of new aggregates is a fibril catalyzed secondary nucleation pathway, that is shared by both the variant of the α-PSMs family (PSMα1 and PSMα3) which sustain the integrity of biofilms [6] and the β-PSM family (PSMβ1). In contrast to this, nucleation and elongation is the only processes contributing to the aggregation of PSMβ2, since the influence of secondary process that give rise to self-replication of aggregates is negligible for this peptide. It is remarkable that even though PSMα1 and PSMα3 possess seven identical and additional 10 similar amino acids in their sequence [10], they show distinct aggregation behavior as PSMα3 aggregates approximately 50 times faster than PSMα1, which specifies that the existence of distinct residue in PSMα3 might play a significant role in lowering the energy barrier for the steps in the conversion process of monomers to fibrils as observed for Aβ peptides [18]. Furthermore the fibrils formed by PSMα3 was found to be initially unstable as also observed before [13] but upon further incubation the fibrils associate laterally to form more mature stable fibrils while fibrils of PSMα1 was stable from without the need for lateral association.

In the current study at quiescent conditions no aggregation kinetics were observed for PSMα4 despite observing β-sheet structure using CD and FTIR and monitoring very thin fibrils with TEM. Previous reports on the aggregation of PSMα4 involved incubation of up to 28 days or incubation under shaking conditions [37, 38]. As shaking conditions during aggregation is known to increase fragmentation due to shear forces [17] shaking conditions were excluded in the present study. Compared to the other PSM peptides PSMα4 is the one with both the lowest solubility score and the lowest calculated aggregation propensity when computing these using the CamSol algorithm [39, 40], Fig S8a-b and Table S3. This could possibly explain the need for long incubation time and shaking conditions during aggregation.

Functional amyloids from gram-negative bacteria are mainly composed of a single protein such as CsgA in *E. coli* curli and FapC in *Pseudomonas* [41, 42]. Along with the proteins incorporated into the functional amyloids a whole range of auxiliary proteins is expressed simultaneously. In the gram-positive bacteria *S. aureus* the functional amyloids in biofilms is made up of the different PSM peptides [6]. The model suggested here account for the role of individual PSM peptides during formation of functional amyloids to stabilize the biofilm hence allowing the bacteria an efficient way to from functional amyloids in a very short amount of time but at the cost of stability. The stability is later gained by the aggregation of other PSM. The most proinflammatory and cytotoxic PSMα3 [12], boosts the reaction of PSMα1 kinetics followed by enhancement of aggregation kinetics of rest of the PSMs peptides, which likely play a key role in stabilizing the biofilm matrix [6] and influences the biofilm development and structuring activities [14]. The highly stable amyloidal structures thus serve as the building blocks cementing the biofilm and creating the rigidity that can explain the resistance of amyloid-containing biofilms. Overall, we note that the rates of individual kinetic steps in the process can differ by several orders of magnitude between different variants, whereas in previous reports a vast structural diversity of amyloid like structures have been also reported for PSMs peptides [37]. Further, *in vitro* studies also confirmed that contrary to what previously thought [6], not all PSMs forms amyloid structures even at higher concentrations.

We conclude that the outcomes presented in this article may have significant implications for understanding the aggregation process of PSMs peptides during biofilm formation. These findings indicate a molecular interplay between individual PSM peptides during accumulation of PSMs amyloid fibrils in biofilms. This also suggest an important approach for suppressing the biofilm growth of *S. aureus* as PSMs have critical role during infection and represent a promising target for anti-staphylococcal activity [43]. Recently potential inhibitors of Aβ aggregation in Alzheimer’s and α-synuclein in Parkinson’s disease have been found to inhibit self-replication by secondary nucleation being the most promising candidate [44]. In the case of *S. aureus* biofilm forming amyloids this could also be a potential strategy as several of the PSM peptides aggregated through a secondary nucleation dominated mechanism. This could be possible by using inhibitors of amyloid formation as numerous studies have demonstrated that inhibitors of aggregation also tend to inhibit biofilm formation [45, 46]. In the context of the development of biofilm formation, the key processes and mechanisms revealed in this study is likely to contribute to the difficulty in controlling and to understanding the role of amyloid growth as a potentially limiting factor of biofilm formation.

## Material and methods

### Peptides and reagents

N-terminally formylated PSM peptides (> 95% purity) were purchased from GenScript Biotech, Netherlands. Thioflavin T (ThT), trifluoroacetic acid (TFA) and hexafluoroisopropanol (HFIP) were purchased from Sigma Aldrich. Dimethyl-sulfoxide (DMSO) was purchased from Merck. Ultra-pure water was used for the entire study.

### Peptide pre-treatment

Lyophilized PSMs peptide stocks were dissolved to a concentration of 0.5 mgml^-1^ in a 1:1 mixture of HFIP and TFA followed by a 5×20 seconds sonication with 30 seconds intervals using a probe sonicator, and incubation at room temperature for one hour. The HFIP/TFA mixture was evaporated by speedvac at 1000 rpm for 3 hour at room temperature. Dried peptide stocks were stored at -80 °C prior to use.

### Preparation of samples for kinetics experiments

All kinetic experiments were performed in 96-well black Corning polystyrene half-area microtiter plates with a non-binding surface incubated at 37 °C in a Fluostar Omega plate reader (BMG Labtech, Germany). Aliquots of purified PSMs were thawed and dissolved in dimethyl sulfoxide (DMSO) to a concentration of 10 mgml^-1^ prior to use. Freshly dissolved peptides were diluted into sterile ddH_2_O containing 0.04 mM ThT. 100 μL of samples were added to each well and the plate was sealed to prevent evaporation. The ThT fluorescence was measured every 10 minutes with an excitation filter of 450 nm and an emission filter of 482 nm at quiescent conditions. However, for PSMα3 the measurement was done every 15 s with same excitation and emission wavelength. The ThT fluorescence was followed by three repeats of each monomer concentration.

### Pre-seeded kinetic assay

Fibrils of different peptides were collected and sonicated for 2 minutes in a sonicator bath at room temperature in low bind Eppendorf tubes (Axygen). Seeds were added to fresh monomer of corresponding peptide immediately before ThT measurements. In cross-seeding experiment, seeds (PSMα1 and α3, PSMβ2) were added to monomer of all other PSMs variants. ThT fluorescence was observed in the plate reader every 10 minutes under quiescent conditions.

### Calculation of the elongation rate constant

To estimate the rates of fibril elongation seeded aggregation with high concentration of preformed fibril seeds (20-50 % of monomeric equivalents concentration) and fixed monomeric concentrations (0.25 mg/mL of PSMα1, 0.5 mg/mL of PSMα3, 0.025 mg/mL for PSMβ1, and 0.25 mg/mL for PSMβ2) was performed. The initial gradients (first 120 min for PSMα1, PSMβ1 and PSMβ2 and the first 120 seconds for PSMα3) of the kinetic curves were determined and plotted against the monomer concentration. Data points at higher concentration were excluded due to saturation effect of elongation.

### Far-UV circular dichroism (CD) spectroscopy

CD was performed on a JASCO-810 Spectrophotometer at 25 °C, wavelength 200 to 250 nm with a step size of 0.5 nm, 2 nm bandwidth and a scan speed of 50 nm/min. Samples were loaded in a 1-mm Quartz cuvette. Triplicate samples containing various peptide concentrations of each freshly dissolved peptides were pelleted and supernatant was transferred to clean sterile tube. The remaining pellet was resuspended in the same volume of ddH_2_O followed by bath sonication. For each sample, the average of five scans were recorded and corrected for baseline contribution and the ddH_2_O signal was subtracted.

### Synchotron radiation circular dichroism spectroscopy (SRCD)

The SRCD spectra of the various PSM fibrils were collected at the AU-CD beamline of the ASTRID2 synchrotron, Aarhus University, Denmark. Three to five successive scans over the wavelength range from 170 to 280?nm were recorded at 25?°C, using a 0.1 mm path length cuvette, at 1?nm intervals with a dwell time of 2 sec. All SRCD spectra were processed and subtracted from their respective averaged baseline (solution containing all components of the sample, except the protein), smoothing with a 7 pt Savitzky-Golay filter, and expressing the final SRCD spectra in mean residual ellipticity. The SRCD spectra of the individual PSM fibrils samples were deconvoluted using DichroWeb [31, 32] to obtain the contribution from individual structural components. Each spectrum was fitted using the analysis programs Selecon3, Contin and CDSSTR with the SP175 reference data set [47] and an average of the structural component contributions from the three analysis programs was used.

### Fourier transform Infrared Spectroscopy (FTIR)

Tensor 27 FTIR spectrometer (Bruker) equipped with attenuated total reflection accessory with a continuous flow of N_2_ gas was used to collect spectra of different aliquots. All samples were dried with N_2_ gas. For each spectrum 64 interferograms were accumulated with a spectral resolution of 2 cm^-1^ in the range from 1000 to 3998 cm^-1^. The data were processed by baseline correction and interfering signals from H_2_O and CO_2_ were removed using the atmospheric compensation filter. Further, peak positions were assigned where the second order derivative had local minima and the intensity was modeled by Gaussian curve fitting using the OPUS 5.5 software. All absorbance spectra were normalized for comparative study.

### Transmission electron microscopy (TEM)

Fibrillated samples (5 μL) of all peptides at various concentrations were placed on carbon coated formvar grid (EM resolutions), incubated for 2 min, washed with ddH_2_O followed by negative staining with 2% uranyl acetate for 2 min. Further, the grids were washed twice with ddH_2_O and blotted dry on filter paper. The samples were examined using a *Morgagni* 268 from FEI Phillips Electron microscopy, equipped with a CCD digital camera, and operated at an accelerating voltage of 80 KV.

### Fibril stability

CD spectra of fibrils (PSMα1, α3, α4, PSMβ1and β2) were recorded from 25 to 95 °C with a step size of 0.1 °C at 220 nm. Stability towards denaturants was tested by dialyzing fibrils (PSMα1, α3, α4, PSMβ1 and β2) containing various concentration of urea (0-8 M) for 24 h.

## Conflict of interest

The authors declare no conflict of interest.

### Acknowledgements

This work was supported by Aarhus University Research Foundation. MA is the recipient of a Starting Grant from Aarhus University Research Foundation. Furthermore we acknowledge the award of beam time on the AU-CD beam line at ASTRID2, under project number ISA-20-1013

## Author contribution

M.Z. and M.A. designed experiments, M.Z. carried out the experiments, M.Z. and M.A. analyzed the data, M.Z. and M.A. wrote the manuscript, M.A. secured the funding.

