## Supplementary material for "Cross-talk between individual phenol soluble modulins in *S. aureus* biofilm formation"

### All together now - Cooperation between individual phenol soluble modulins in S. aureus biofilm formation

#### Molecular mechanism of functional amyloid formation in *S. aureus* biofilms

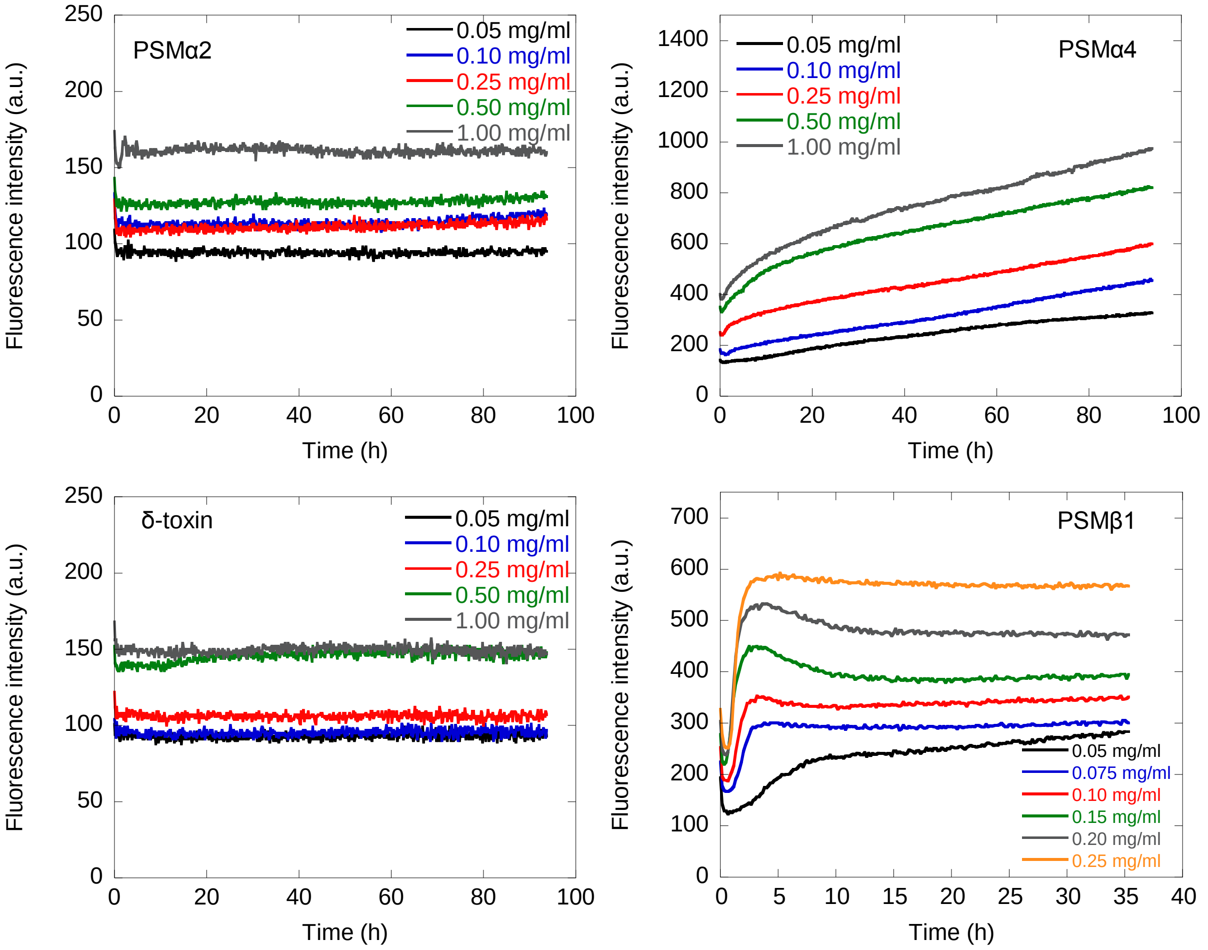


**Figure S1:** Experimental kinetic data for the aggregation of the PSMs peptides under quiescent conditions. Aggregation of PSMs peptides (a) PSMα2, (b) PSMα4, (c) δ-toxin and (d) PSMβ1from monomeric samples (0.05 mg/ml to 1.0 mg/ml) is measured by ThT fluorescence at 37°C every 10 min. Three repeats were carried out at each condition. At higher concentrations of βPSM1 the lag-time becomes independent of the monomer concentration indicating some sort of saturation effect.

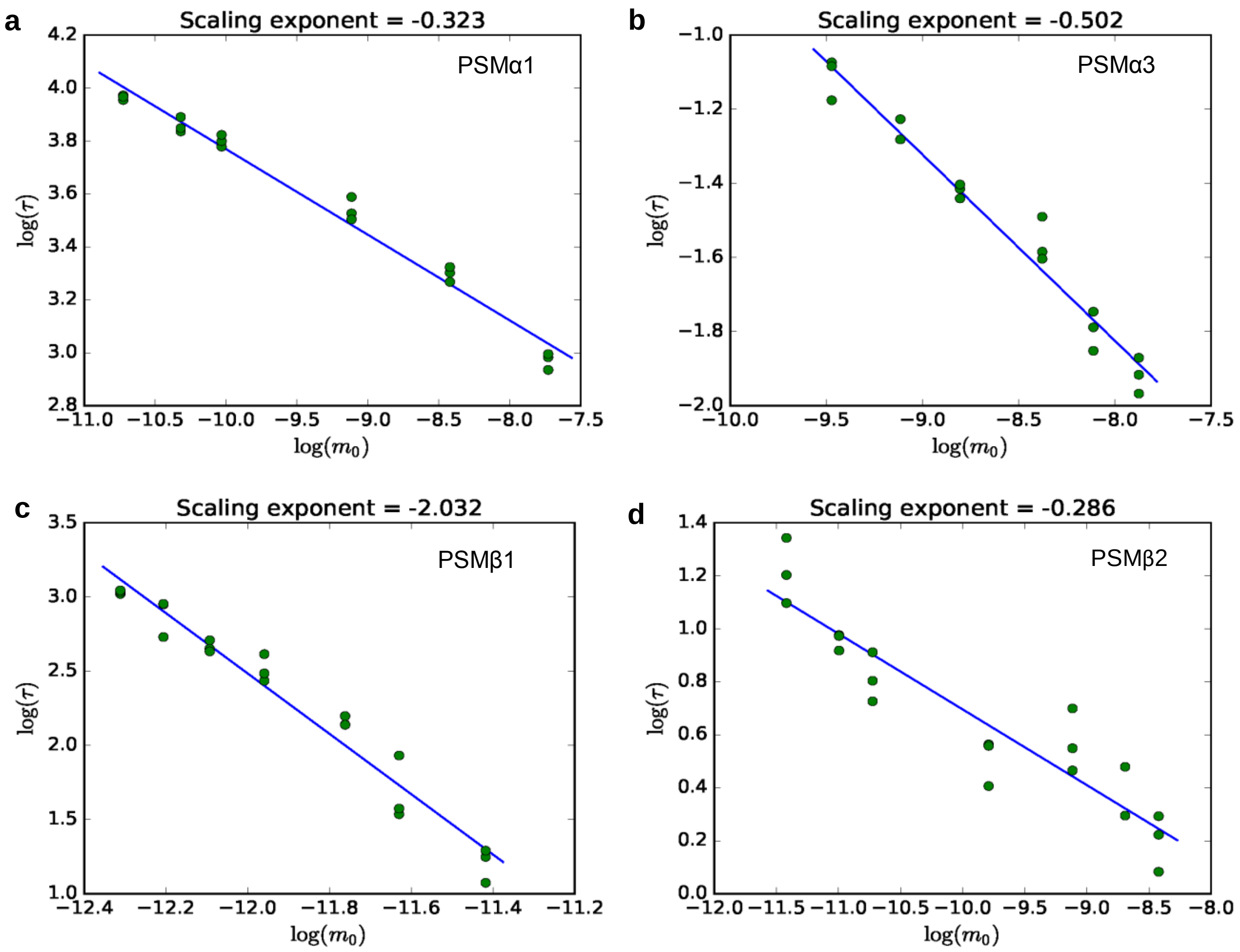


**Figure S2**: The half times of fibril formation as a function of initial monomer concentration on double logarithmic axes. The slope of the fitted line gives the scaling exponent (γ), obtained from three repeats of the aggregation experiments for each peptide. (a) PSMα1, (b) PSMα3, (c) PSMβ1, and (d) PSMβ2. The straight line in all three plots indicates that the dominant mechanism of fibril multiplication is the same for all monomer concentrations for the individual peptides studied here.

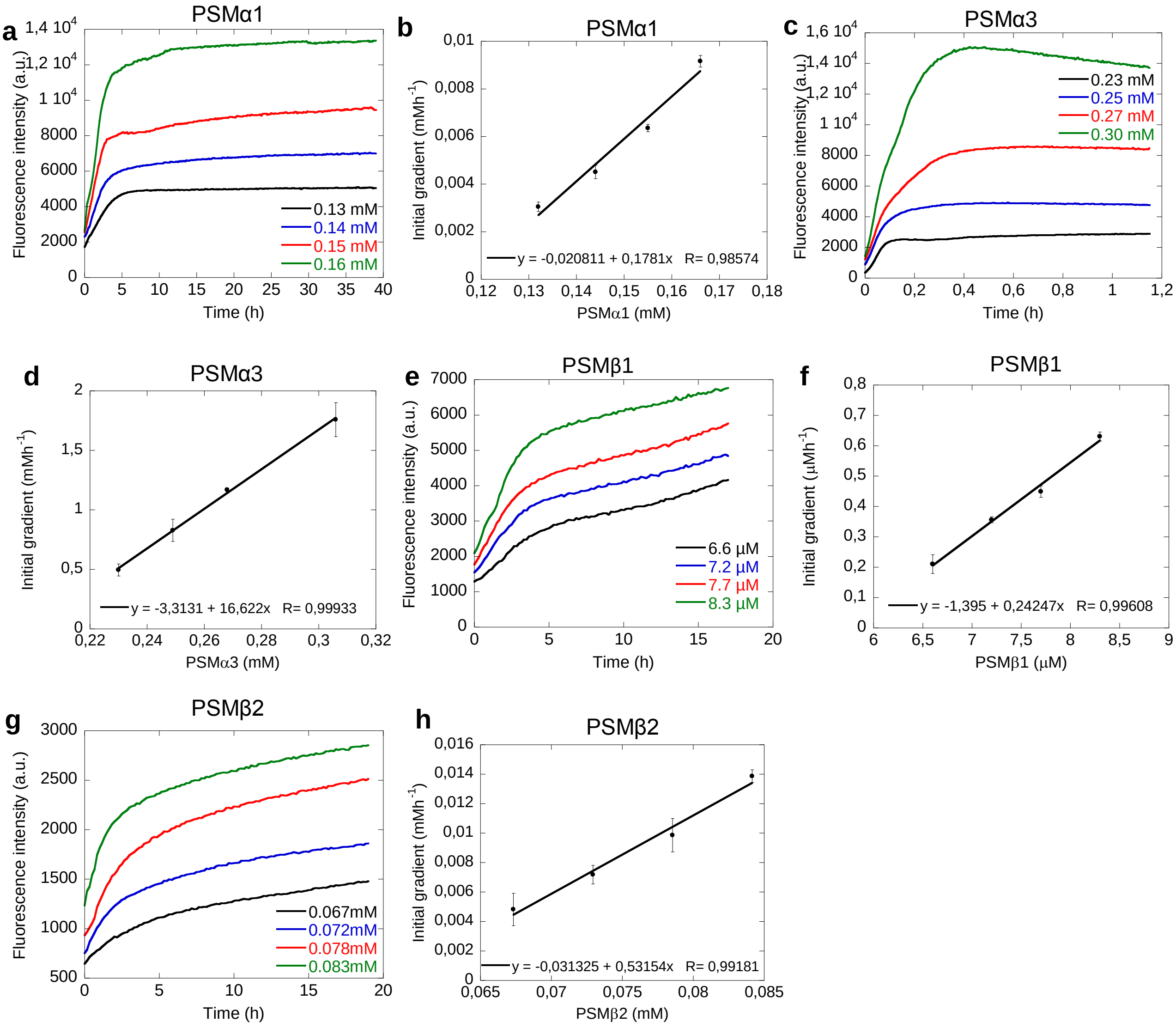


**Figure S3:** Seeing of PSM peptides with 20-50 % preformed fibrils seeds (in monomeric equivalents) at different monomeric concentration. (a) Changes in ThT fluorescence when monomeric αPSM1 at fixed concentration (0.25 mg/mL) was incubated in presence of high concentrations of preformed fibrils from the αPSM1 under quiescent conditions at 37°C. (b) Initial gradient of the ThT fluorescence curves of PSMα1used to estimate the elongation rates. The initial gradient (first 120 min) is plotted against the free monomer concentration. A straight line was fitted to these points, with the slope proportional to the number of seed fibrils and the elongation rate constant. (c) Changes in ThT fluorescence when monomeric αPSM3 at fixed concentration (0.5 mg/mL) was incubated in presence of high concentrations of preformed fibrils from the αPSM3 under quiescent conditions at 37°C. (d) Initial gradient of the ThT fluorescence curves of PSMα3 used to estimate the elongation rates. The initial gradient (first 2 min) is plotted against the free monomer concentration. A straight line was fitted to these points, with the slope proportional to the number of seed fibrils and the elongation rate constant. (e) Changes in ThT fluorescence when monomeric βPSM1 at fixed concentration (0.025 mg/mL) was incubated in presence of high concentrations of preformed fibrils from the βPSM1 under quiescent conditions at 37°C. (f) Initial gradient of the ThT fluorescence curves of PSMβ1used to estimate the elongation rates. The initial gradient (first 120 min) is plotted against the free monomer concentration. A straight line was fitted to these points, with the slope proportional to the number of seed fibrils and the elongation rate constant. (g) Changes in ThT fluorescence when monomeric βPSM2 at fixed concentration (0.25 mg/mL) was incubated in presence of high concentrations of preformed fibrils the βPSM2 under quiescent conditions at 37°C. (b) Initial gradient of the ThT fluorescence curves of PSMβ2 used to estimate the elongation rates. The initial gradient (first 120 min) is plotted against the free monomer concentration. A straight line was fitted to these points, with the slope proportional to the number of seed fibrils and the elongation rate constant.

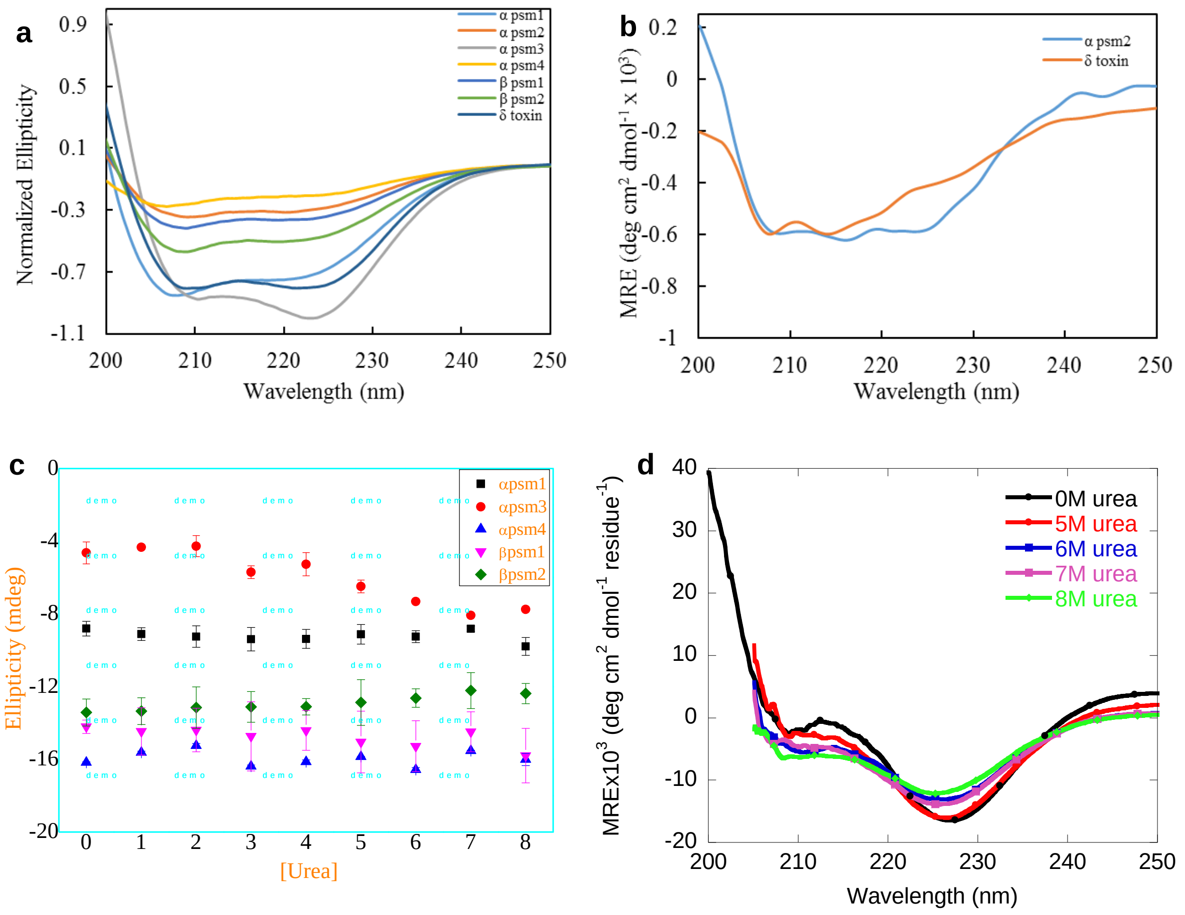


**Figure S4:** CD spectra of monomeric PSM peptides and of PSM peptides that do not aggregate. (a) CD spectra of monomeric PSMs peptides (0.25 mg/ml) measured at 25°C followed by 0 h of incubation period. (b) Far UV-CD spectra of PSMα2 and δ-toxin after 7 days incubation at 37C display α-helical character consistent with the lack of aggregation. (c) Analysis of the dissociation of PSMs fibrils after incubation in presence of various concentrations (0-8M) of urea. (d) Far UV-CD spectra of PSMα3 fibrils after incubation in the absence and presence of high concentrations of urea (0 and 5-8 M urea).

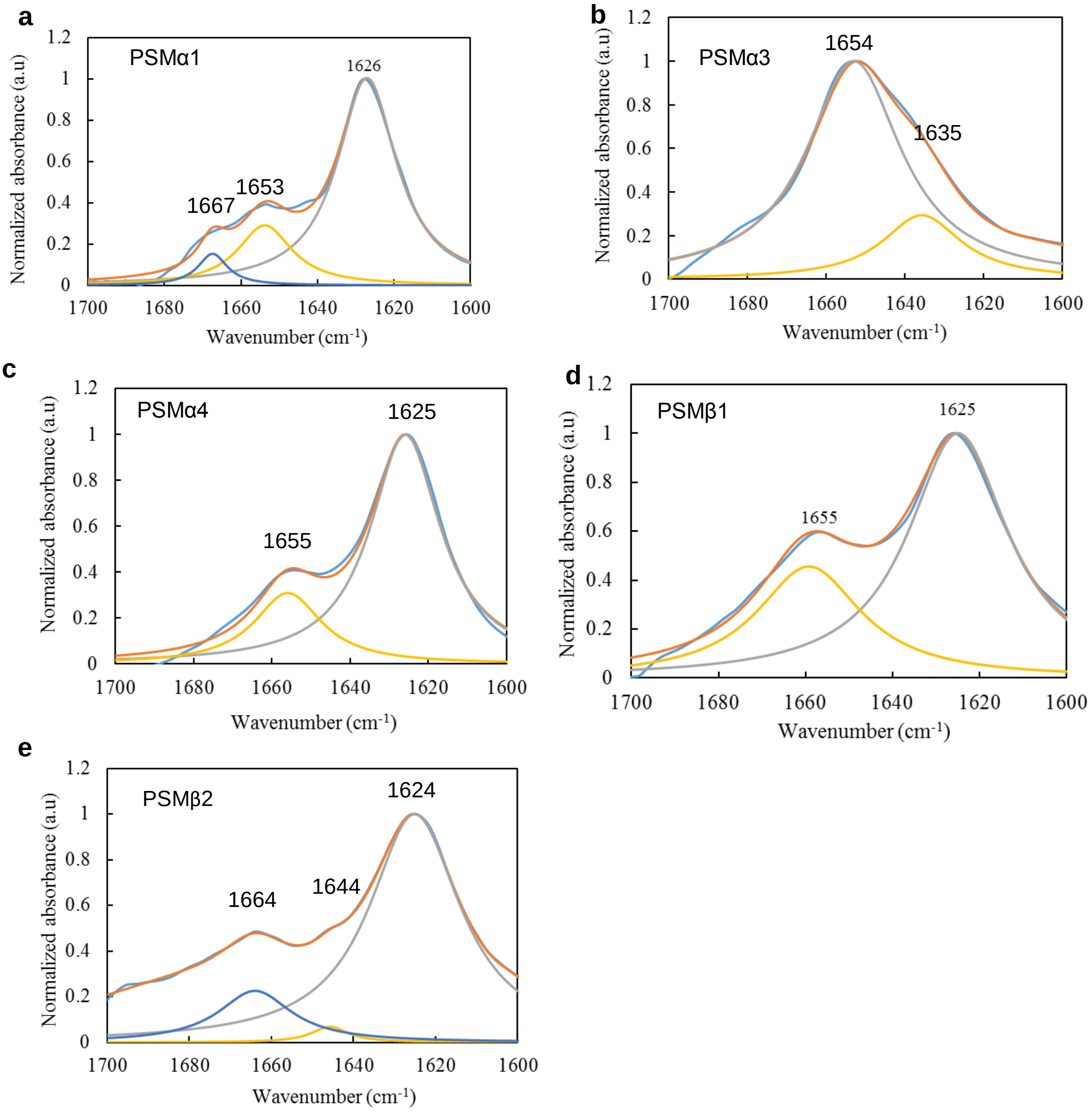


**Figure S5:** FTIR analysis of the secondary structure of PSMs peptides. (a) FTIR spectra and second order derivative of the spectra of PSMα1 at 0.5 mg/mL. (b) FTIR spectra and second order derivative of the spectra of PSMα3 at 0.5 mg/mL. (c) FTIR spectra and second order derivative of the spectra of PSMα4 at 0.5 mg/mL. (d) FTIR spectra and second order derivative of the spectra of PSMβ1 at 0.25 mg/mL. (e) FTIR spectra and second order derivative of the spectra of PSMβ2 at 0.5 mg/mL.

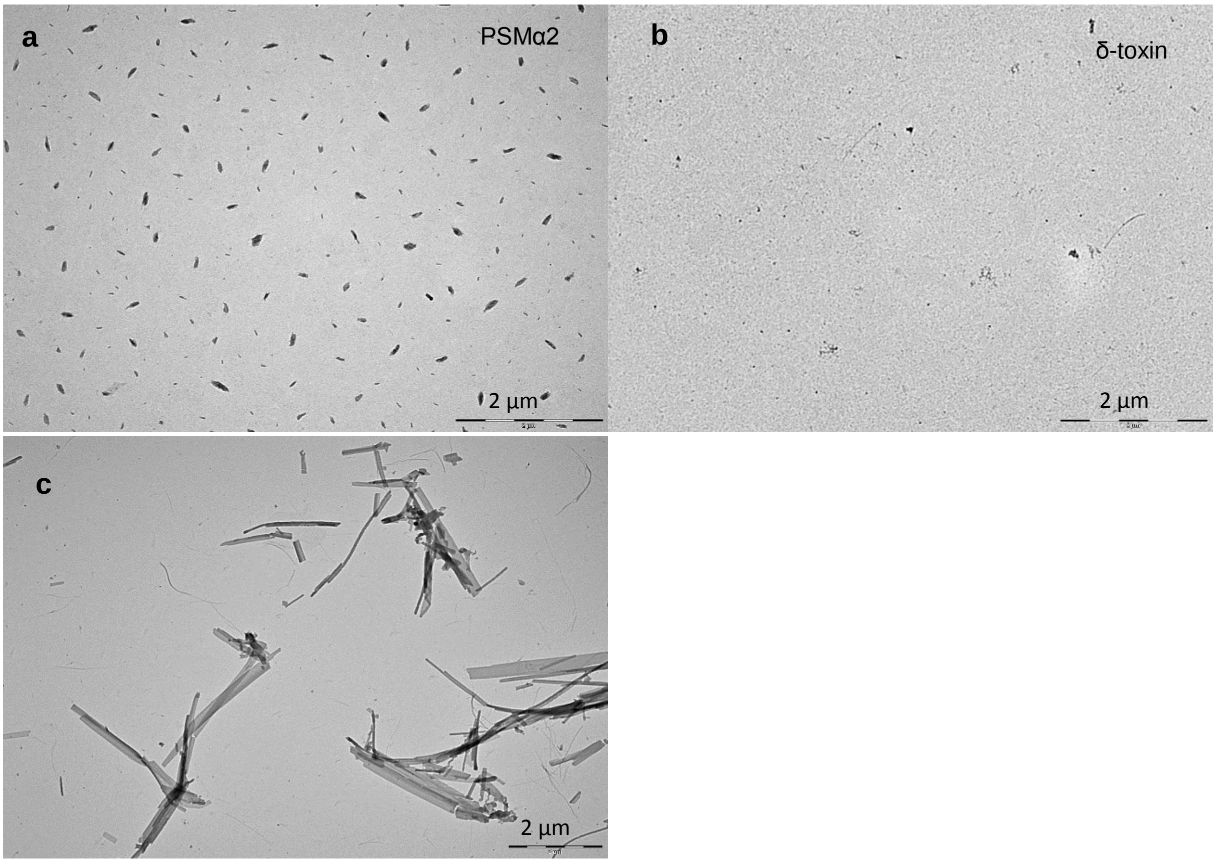


**Figure S6:** Morphology of PSMs peptides. Transmission electron microscopic image of the end state of reaction for samples initially composed of (a) PSMα2 monomers, (b) δ-toxin monomers and (c) PSMα3 after two days of incubation.

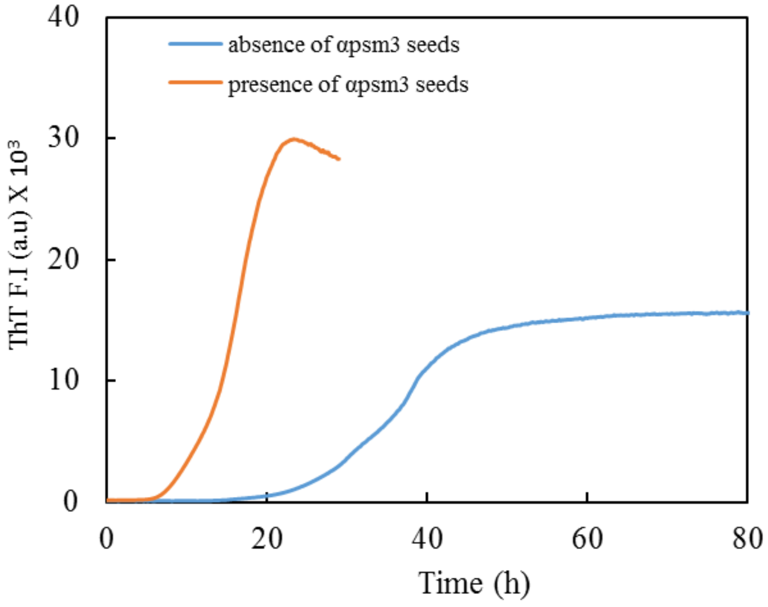


**Figure S7:** The acceleration of the aggreagtion of PSMα1 upon addition of PSMα3 preformed fibril seeds to monomeric PSMα1 peptide.

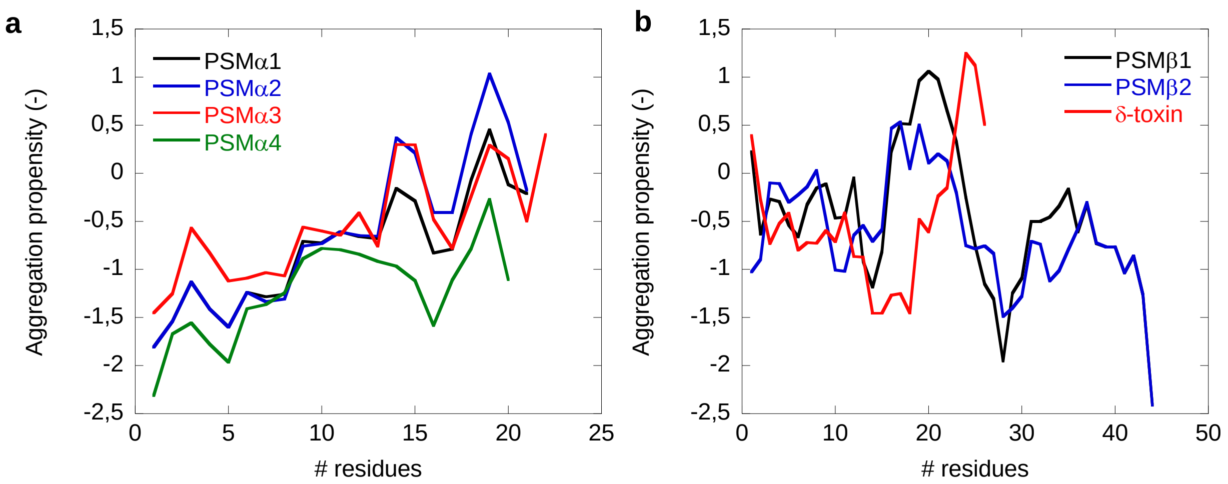


**Figure 8:** Aggregation propensity profiles of PSM peptides using CamSol [38, 39]. (a) Aggregation propensities of PSMα group of peptides (PSMα1, PSMα2, PSMα3 and PSMα4). (b) Aggregation propensity of PSMβ group of peptides (PSMβ1 and PSMβ2) along with δ-toxin.

**Table S1:** Percentage contribution of various structural components for the fibrils of PSM variants based on deconvolution of SRCD spectra using the DichroWeb server using the reference data setSP175 for the Selecon3, Contin and CDSSTR analysis programs .

| Peptide | α-helix | β-sheet | Turns | Unordered |
|---|---|---|---|---|
| PSMα1 | 16,1 | 33,1 | 10,5 | 38,5 |
| PSMα3 | 69,3 | 11,3 | 6,1 | 13,2 |
| PSMα4 | 23,4 | 31,0 | 10,1 | 35,4 |
| PSMβ1 | 34,5 | 23,8 | 10,1 | 33,0 |
| PSMβ2 | 39,5 | 16,4 | 10,6 | 34,7 |

**Table S2:** Percentage contribution of various structural components for the fibrils of PSM variants based on deconvolution of FTIR spectra along with peak position.

| Peptide | Peak position | β-sheet | α-helix | β turns |
|---|---|---|---|---|
| PSMα1 | 1626, 1653,1667 | 68.58 | 26.32 | 9.10 |
| PSMα3 | 1635, 1654 | 43.94 | 56.06 | - |
| PSMα4 | 1625, 1655 | 69.12 | 30.88 | - |
| PSMβ1 | 1625, 1656 | 66.80 | 33.2 | - |
| PSMβ2 | 1624,1644, 1664 | 66.22 | 20.05 | 13.73 |

**Table S3:** Solubility score of different peptides calculated using the Camsol web server (http://www-mvsoftware.ch.cam.ac.uk/index.php/login):

| Peptide Name | Solubility Score |
|---|---|
| PSMα1 | 0.826149 |
| PSMα2 | 0.883944 |
| PSMα3 | 1.282062 |
| PSMα4 | 0.022660 |
| PSMβ1 | 1.038055 |
| PSMβ2 | 0.907007 |
| δ-toxin | 1.242128 |
